# Group 2 Innate lymphoid cells promote allograft survival by constraining and inducing anergy in alloreactive CD4^+^ T cells

**DOI:** 10.1101/2025.06.03.657622

**Authors:** Jifu Ge, Weikang Pan, Zijie Xu, Thais Boccia da Costa, James F. Markmann, Ivan Zanoni, Alex G. Cuenca

## Abstract

Although solid organ transplant outcomes have dramatically improved over the last several decades, incomplete understanding of the immune interface between the donor organ and the recipient’s immune system has impaired our ability to induce immune tolerance in most transplant recipients. Since group 2 innate lymphoid cells (ILC2s) reside in all transplanted solid organs, participate in wound healing, and coordinate other immunoregulatory cell populations, we investigated their role in the alloimmune response. Using a mouse heterotopic cardiac transplant model, we show that recipient ILC2s replace donor’s ILC2s, upregulate MHCII without expressing costimulatory molecules. In addition, recipient derived ILC2s process and present alloantigen, inducing CD4^+^ T cell anergy via Caspase-3 pathway. When recipient-derived ILC2s are not present, we observed a significant increase in infiltrating donor reactive CD4^+^ T cells and worsened allograft survival. Additionally, expansion of ILC2s in vivo through IL33 administration prolonged the survival of murine heart allografts. Overall, these data highlight a critical and novel immunoregulatory role of host-derived ILC2s in solid organ transplant, where they induce anergy in alloreactive CD4^+^ T cells, promoting the induction of alloimmune tolerance.

## Introduction

Solid organ transplantation remains an invaluable treatment for end-stage organ failure. Despite advances in surgical technique, organ preservation, and immunosuppression, morbidity associated with deleterious but necessary medications remain a major challenge^1,2^. A significant effort has been made towards the development of cellular therapeutics that decrease or obviate the need for these immunosuppression regimens. These interventions have mainly focused on the adaptive immune response^3^. The role of innate immunity in alloimmune responses post-transplant remains unclear.

Innate lymphoid cells (ILCs) are tissue resident innate immune effector cells that are divided into three subsets based on the expression of the transcription factors T-bet (ILC1), GATA3 (ILC2), and RORγt (ILC3)^4^. ILCs rapidly respond to environmental signals and tissue damage, influencing immune responses through cytokine release and interactions with other cells. In addition, there is a growing body of literature to suggest that ILCs also play an important role in transplant related immunity. In lung transplant recipients, primary graft dysfunction (PGD) was associated with a decrease in the ILC2 subset following reperfusion^5^. In contrast, individuals who did not develop PGD presented with increased ILC1 levels before reperfusion, followed by higher ILC2 levels after the allograft reperfusion. ILC2s have also been shown to alleviate acute gastrointestinal GVHD when adoptively transferred to HSCT recipients in a murine study^6^.

Herein, we investigated the role of ILC2s during mouse heart transplantation. We reveal that recipient ILC2s infiltrate transplanted hearts and upregulate the expression of MHCII in the absence of CD80/CD86. In addition, we show that ILC2s are capable of processing and presenting alloantigens within murine cardiac allografts. We also demonstrate that infiltrating recipient ILC2s, which lack costimulatory molecules, induce apoptosis and anergy of CD4^+^ T cells in an MHCII dependent manner in vitro. In vivo, ILC2s constrain alloimmune T cell infiltration and support allograft survival. Overall, we identified a novel mechanism by which ILC2s contribute to alloimmune responses following solid organ transplantation, paving the way to therapeutically manipulate ILC2s to provide allograft tolerance.

## Results

### Recipient derived ILC2s replenish donor’s ILC2s in cardiac allografts following heart transplant

ILC2s are tissue resident cells in both lymphoid and non-lymphoid organs that were initially thought to be maintained in a self-renewing manner during both physiological and inflammatory conditions^7^. However, ILC2s also have the capacity to migrate and restore tissue-resident ILC2 populations^8^. How donor and recipient ILC2 populations change following heart transplant remains unknown. To examine this, we examined the presence of ILC1, ILC2 and ILC3 populations in the hearts of C57BL/6 (B6) mice under homeostasis and following murine heterotopic cardiac transplant (mHTx). As previously reported, Lineage(Lin)-CD127^+^GATA3^+^ ILC2s represent the largest ILC population in the naïve heart, with less than 5% of ILC1s or ILC3s detected in the Lin-CD127^+^ population (Figure 1A and 1B)^4,10^. Cardiac CD127+GATA3+ ILC2 populations expressed high levels of ST2, CD90, KLRG1 and CD25 (Figure 1C). Similarly, ILC1s or ILC3s were rarely found in cardiac allografts of Balb/C (H2-Kd) to B6 (H2-Kb), an MHC I and II fully mismatched combination, post operative day 7 following mHTx (Figure 1D).

**FIGURE 1.**
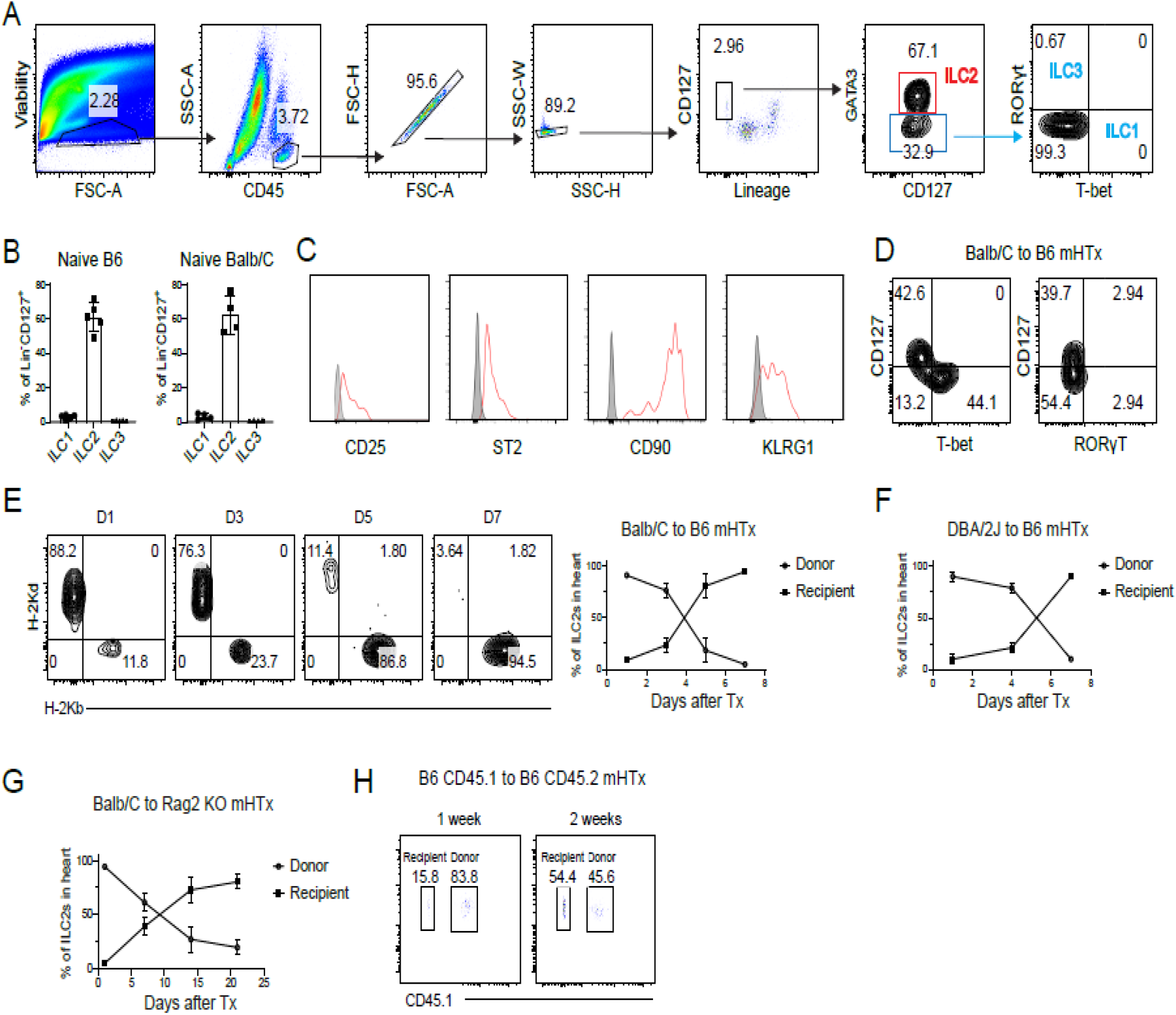
Recipient ILC2s migrate into heart allografts following transplant and replace donor ILC2s. Single-cell suspensions from naïve hearts and grafts from murine heart transplants (mHTx) were prepared to assess ILC composition and changes in ILC2 populations following heart mHTx. (A) Representative flow cytometry plots showing the gating strategy for characterizing ILC populations in heart tissue. CD45^+^Lineage (Lin)-CD127^+^ cells were identified as ILCs. ILC subsets were defined as follows: ILC2 (Lin-CD127^+^GATA3^+^), ILC1 (Lin-CD127^+^T-bet^+^), and ILC3 (Lin-CD127^+^RORγt^+^). (B) Bar graphs showing the relative proportions of ILC1, ILC2, and ILC3 cells within the Lineage-CD127^+^ population in naïve heart. Data are presented as the percentage of each ILC subset within the total Lin-CD127^+^ population (n=5 mice/group). (C) Representative histograms showing surface markers for ILC2s. (D) Representative flow cytometry plots of ILC1 (CD127^+^T-bet^+^) and ILC3 (CD127^+^RORγt^+^) populations. Single-cell suspensions from cardiac allografts following Balb/C to B6 heart transplants were gated on the CD45^+^ Lin-population. (E - G) Flow cytometry scatter plots showing the changes of ILC2 populations derived from both donor hearts (H2-Kd) and recipient(H2-Kd) at indicated time points following transplantation. Line graphs depict the changes in the proportion of donor- and recipient-derived ILC2s across different transplant combinations as indicated over time (n=5/group). (H) Flow cytometric plots of donor and recipient ILC2s in graft following syngenetic pep-boy B6(CD45.1) to B6(CD5.2) hearts transplant. ILC2s were isolated from cardiac grafts and examined for donor (CD45.1) and recipient (CD45.2) derived ILC2s.

Gasteiger et al reported that heart resident ILC2s renew locally within tissue with very few cells replenished from hematogenous sources, as parabiosis experiments showed limited numbers of ILC2s circulate in-between parabiotic partners^7^. We sought to understand whether recipient derived ILC2s replenish the donor heart graft ILC2s cell pool after mHTx. An active migration of recipient derived ILC2s to the donor heart grafts was observed in this acute rejection model (Figure 1E). Similar results were obtained using Balb/C congenic strain, DBA/2J (H2-Kd) to B6 mHTx, suggesting that the replenishing of the donor ILC2 pool with recipient-derived ILC2s is not strain specific (Figure 1F).

To understand whether the reconstitution of cardiac ILC2s relies on the adaptive immune system, we used Rag2 KO mice which lack mature T cells and B cells as recipients for heart transplant. Although in a slower manner, recipient-derived ILC2s still migrated into donor hearts allowing 90% replacement of donor ILC2s with recipient ILC2s occurring at 3 weeks (Figure 1G). To understand if this process occurs due to alloimmune recognition or deletion, we used hearts from B6 syngeneic pep-boy donors (CD45.1^+^) and transplanted them into CD45.2^+^ B6 recipients. Recipient ILC2s (CD45.2^+^) gradually replaced donor ILC2s (CD45.1^+^), approximately half of the heart ILC2s were reconstituted by recipient-derived ILC2s by 2 weeks (Figure 1H). These data suggest that, similar to studies by Cautivo or Shaikh, in pathological or inflammatory circumstances, mHTx leads to migration and replacement of donor-derived ILC2s with recipient ILC2s^8,9^.

### Graft infiltrating ILC2 upregulate histocompatibility complex class II associated pathways

ILC2 subsets acquire different transcriptional programs depending on the tissue they reside^8^. To study this, we isolated RNA from ILC2s sorted from naïve hearts or cardiac allografts post-operative day (POD) 7 following mHTx. Differential gene expression analysis was performed on these two groups of samples using Smart-Seq data. A total of 701 differentially expressed genes (DEGs) were identified, with 381 upregulated and 320 downregulated genes (adjusted p-value < 0.05, |log2 fold change| > 0.586). The distribution of DEGs is illustrated in a volcano plot (Figure 2A), highlighting a significant enrichment of genes associated with major histocompatibility complex class II (MHCII) pathways.

**FIGURE 2.**
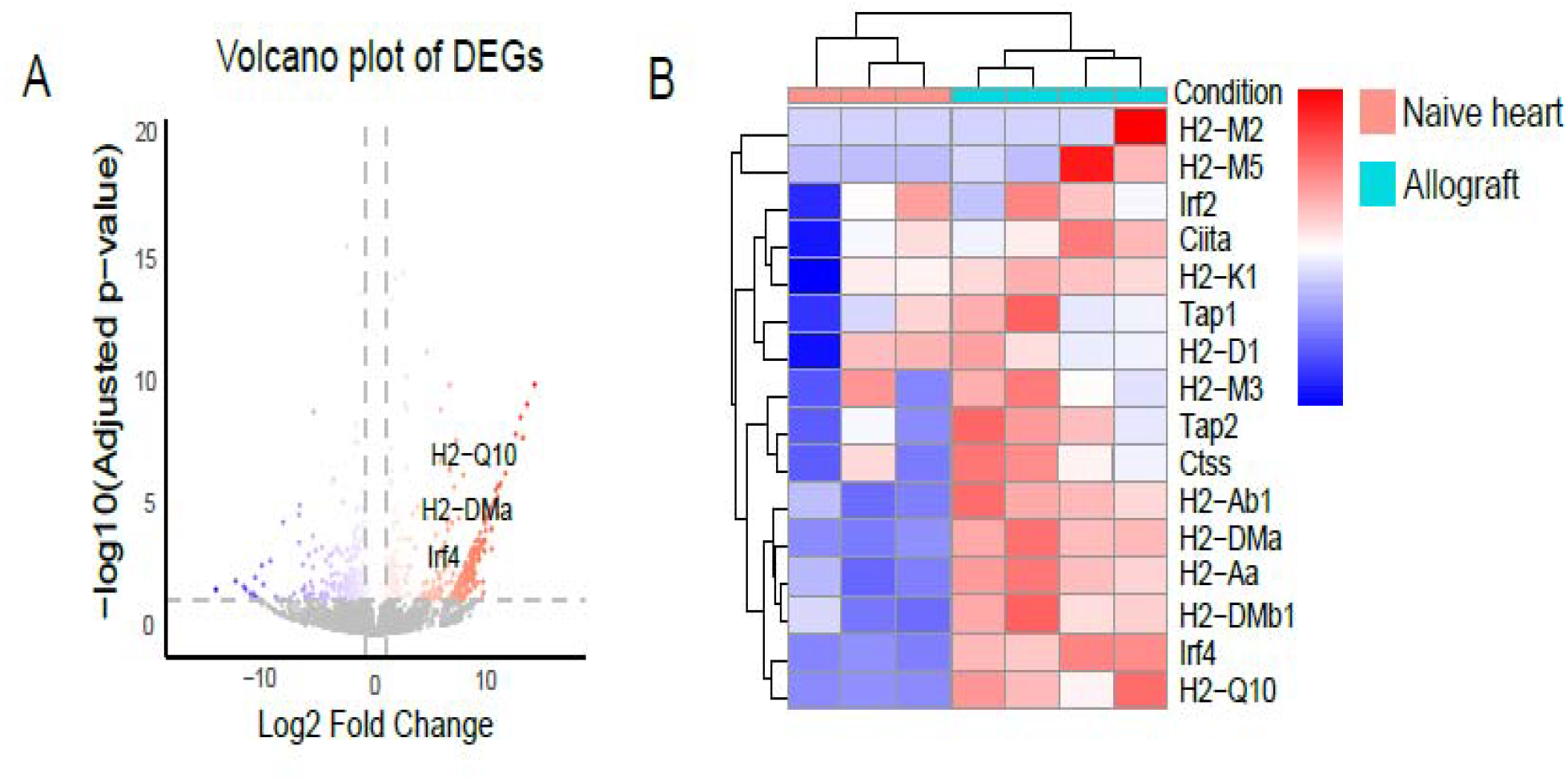
Smart-Seq analysis reveals allograft ILC2s are transcriptionally distinct and enriched for MHCII related genes post heart transplant. (A) Volcano plot showing differentially expressed genes (DEGs) between naïve and graft-infiltrating ILC2s. Red dots represent genes with statistically significant differential expression, identified using a false discovery rate (FDR) < 0.05 and a minimum 2-fold change in expression (log2FC > 1). The plot highlights the upregulation of several genes associated with antigen presentation in graft-infiltrating ILC2s. (B) Heatmap comparing genes associated with antigen presentation in naïve versus POD7 cardiac allograft ILC2s. To obtain sufficient RNA, ILC2s from naïve hearts were pooled from 4-7 naïve hearts to generate a single sample, with a total of n=3 replicates used for comparison with ILC2s from cardiac allografts.

To further investigate the expression patterns of these DEGs, a heatmap was generated (Figure 2B), showing distinct clustering of MHCII-related genes between the two groups. Key upregulated MHCII-related genes included *Ciita, Ctss, H2-Aa and H2-Ab1*, etc. These results suggest that the differential regulation of MHCII genes may play a critical role in the biological processes distinguishing the two groups.

### Graft Infiltrating ILC2s upregulate MHC II without expressing co-stimulatory molecules CD80/86

ILC2s have been shown to facilitate wound repair, interact with T helper cells in a paracrine manner, and have intrinsic immunoregulatory properties^11,12^. ILC2s can also potentially establish direct interactions with alloimmune T cells. To study this, we first analyzed whether MHCII is constitutively expressed on ILC2s at baseline. Several investigators have reported that ILC2s can upregulate MHCII upon parasitic infection^12,13^. Indeed, we detected a small proportion of ILC2s with the MHCII in the naïve heart (Figure 3A). Following transplant, however, ILC2s infiltrating the graft significantly upregulated the expression of MHCII at 1, 3, and 6 weeks post mHTx in Balb/C into Rag2 KO mice and at 3, 5, and 7 days in Balb/C into B6 transplants (Figure 3A and 3B). To further delineate whether MHCII expression was present on either donor and/or recipient derived ILC2s, we assayed for the expression of MHCII on H2-Kb positive versus negative cells (Figure 3C). As demonstrated in Figure 3C, compared to donor ILC2s, recipient ILC2s significantly upregulated MHCII following transplant in either Balb/C into Rag2 KO or the Balb/C into B6 model.

**FIGURE 3.**
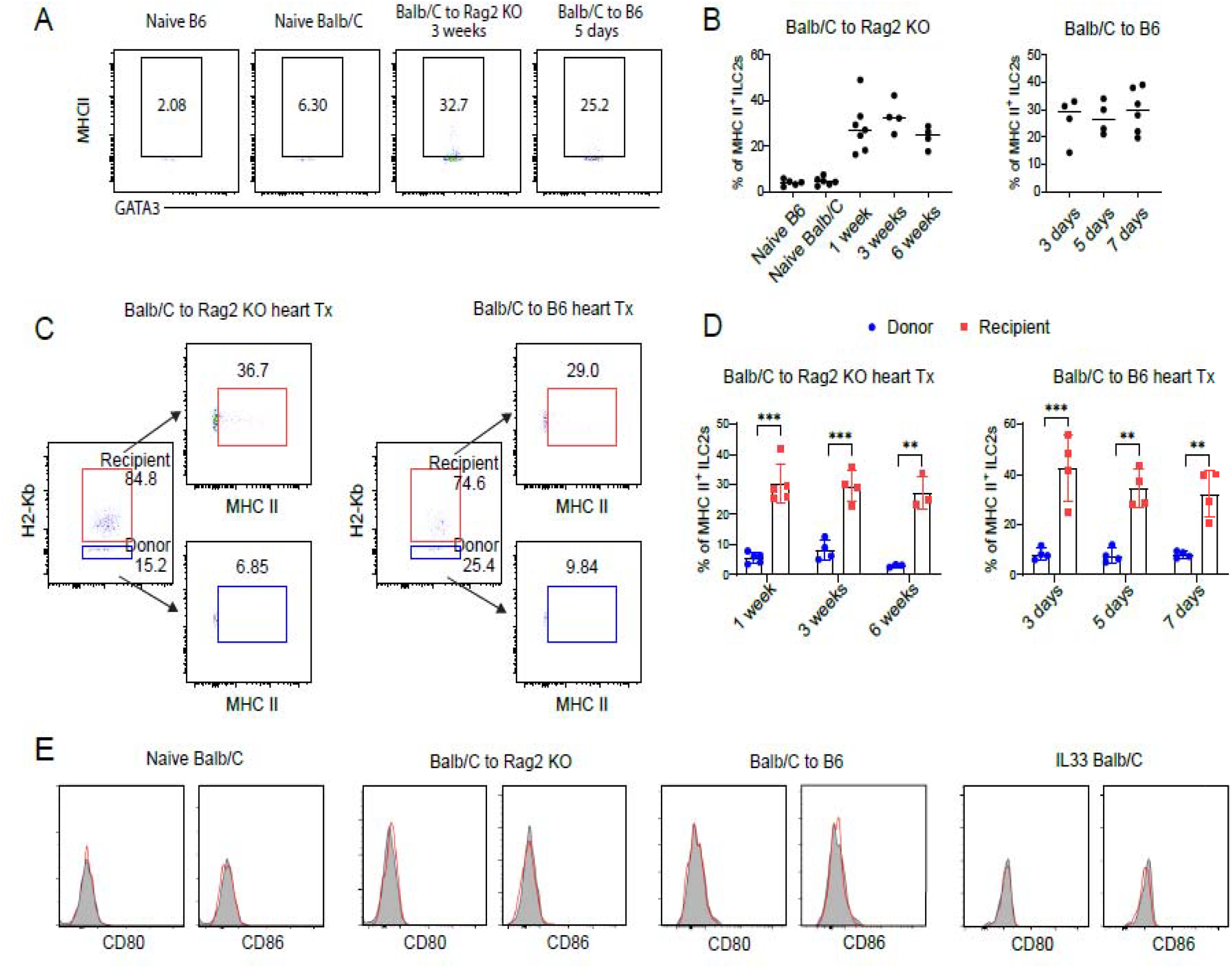
Recipient ILC2s upregulate MHCII following solid organ transplant. (A and B) Representative flow cytometric plots and statistical analysis of MHCII expression on ILC2s from naïve hearts or cardiac allografts at different time points post murine heart transplant as indicated. MHCII expression was assayed via flow cytometry on ILC2s in both an immunocompromised (Balb/C to Rag2 KO) and immunocompetent (Balb/C to B6) model (n=4-5/group). (C and D) Representative flow cytometry figures and statistic analysis of MHCII expression on donor and recipient derived ILC2s in cardiac allografts. ILC2s from allografts were labeled for H2-Kb to identify donor vs recipient populations, and assayed for MHCII expression at different time points after transplant (n=4-5; ** p< 0.01 *** p< 0.001). (E) Representative flow cytometric histogram figures showing expression of CD80/CD86 on heart ILCs. ILC2s were isolated from heart of naïve or IL-33 conditioned mice, and allografts from transplant recipients as indicated, and assayed for the presence of CD80/86 on the cell surface of ILC2s.

Antigen-presenting cells (APCs) process and present antigens via MHCII and interact with CD4^+^ T cells, potentially activating them or inducing anergy, depending on the expression of co-stimulatory molecules by the APCs. To determine whether graft-infiltrating ILC2s express CD80/CD86 and their potential impact on CD4^+^ T cells, single cell suspensions from naïve murine hearts as well as cardiac allografts 7 days post mHTx were examined for CD80 and CD86 expression. CD80 or CD86 were not expressed on ILC2s from naïve murine hearts or heart allografts, while ILC2s from mesenteric lymph nodes (mLNs), lung, and kidney expressed these co-stimulatory molecules constitutively. Although ILC2s from mLNs and lung, and to a lesser extent kidney, showed significant upregulation of CD80 and CD86 following IL33 activation *in vivo*, expression of these molecules on heart ILC2s from IL33-treated mice remained low and did not differ significantly from baseline (Supplemental Figure 1 and Figure 3D). These findings highlight the tissue-specific regulation of co-stimulatory molecule expression on ILC2s, with significant differences between heart from various other tissues, including mesenteric lymph nodes, lung, and kidney. This suggests that ILC2s may exhibit distinct immunoregulatory properties depending on their tissue microenvironment, leading to the hypothesis that interaction of ILC2s with alloreactive CD4^+^ T cells in the newly transplanted solid organ could lead to CD4^+^ T cell anergy.

### Presence of MHCII on ILC2s following heart allograft transplant is not related to cross-dressing

Professional antigen presenting cells (APCs) such as dendritic cells, B cells, and macrophages, have been shown to acquire MHCII molecules or MHCII-peptide complex from other cells through trogocytosis, exosomes and nanotubes, a process termed “cross-dressing”^14^. To address whether ILC2s infiltrating the cardiac allograft undergo cross-dressing, we assayed recipient derived ILC2s (H2-Kb) isolated from heart grafts for the presence of donor MHCII (Balb/C, I-Ad). However, no donor MHCII (I-Ad) was detected on recipient ILC2s (H2-Kb) after heart transplant (Figure 4A). As immunocompetent B6 mice reject Balb/C mouse hearts at approximately 8 days post-mHTx, it is possible that ILC2s could obtain donor MHCII at later time points through interaction with recipient APCs. To address the possibility that cross-dressing in our model requires longer ILC2-donor APC interactions, we used Rag2 KO mice on B6 background as recipients, which take significantly longer to reject heart grafts. Similar to data in Figure 4A, recipient-derived ILC2s upregulated MHCII upon transplant 21 days post heart transplant, while no donor MHCII I-Ad was found on ILC2s from allograft recipients (Supplemental Figure 2). We did identify Rag2 KO recipient derived lineage positive populations (Lin^+^H2-Kb^+^, consisting of recipient professional APCs including DCs and macrophages) that expressed significant amounts of donor MHCII I-Ad, suggesting that active cross dressing occurred on other cell types within the cardiac allograft but not ILC2s (Figure 4B). Moreover, ILC2s isolated from MHCII KO mice did not show MHCII expression after adoptively transferred into Rag2/Il2rg DKO mice that received Balb/C heart, further supporting that ILC2s do not receive cell surface MHCII from other cells post-transplant via cross-dressing (Figure 4C).

**FIGURE 4.**
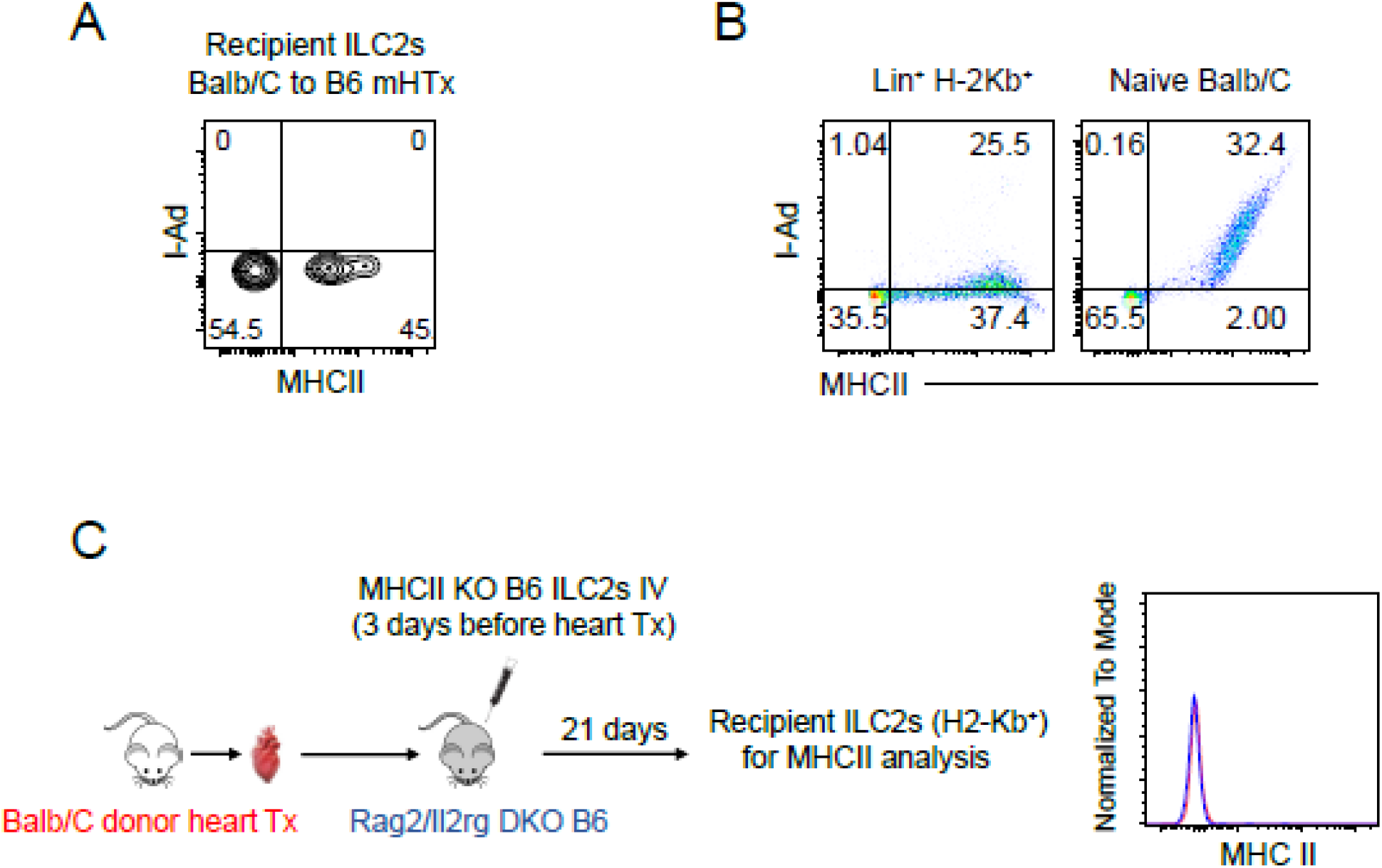
MHCII expression on ILC2s following heart transplantation is not due to cross-dressing. (A) Flow cytometry analysis of recipient ILC2s (H2-Kb^+^) from B6 mice receiving Balb/C heart allografts. Plots show MHCII (I-Ab) vs. donor MHCII (I-Ad) expression. (B) Flow cytometry analysis of Lin^+^ H2-Kb^+^ cells (left) and naive Balb/C cells (right) for donor MHCII (I-Ad) expression. Naïve Balb/c hearts immune cells were assayed for MHCII (I-Ad) as positive control (C) Left: Experimental schematic showing MHCII KO B6 ILC2s intravenously injected into Rag2/Il2rg DKO B6 recipients 3 days before Balb/C heart transplantation. Analysis was performed 21 days post-transplant. Right: Representative histogram showing MHCII expression on recipient ILC2s (H2-Kb^+^) isolated from heart allografts.

### Graft infiltrating ILC2s can process and present donor antigen

Although we found that graft-infiltrating ILC2s can upregulate MHCII, it remains unclear whether they can process and present antigens like professional APCs. To investigate whether ILC2s possess the ability to endocytose and process antigens, soluble protein DQ-ovalbumin (DQ-OVA) which produce bright fluorescence during hydrolysis by proteases, was cultured with heart ILC2s in vitro. Cardiac derived ILC2s exhibited a strong fluorescent signal as early as 2 hours after culturing with DQ-OVA, indicating that they can uptake and degrade soluble antigen (Figure 5A)^12^. Next, we determined whether infiltrating ILC2s could not only process antigen but also present alloantigens. To understand this, we isolated ILC2s from immunocompetent mHTx recipients at day 7 post mHTx and stained for MHCII-antigen complex. As shown in Figure 5B, we found the cell surface expression of donor peptide I-Eα_52-68_ (Balb/C) in the context of B6 derived MHCII (I-Ab) on recipient ILC2s by using the antibody Y-Ae ^15^. These data show that recipient ILC2s migrating to the allograft can both process and present donor derived antigen.

**FIGURE 5.**
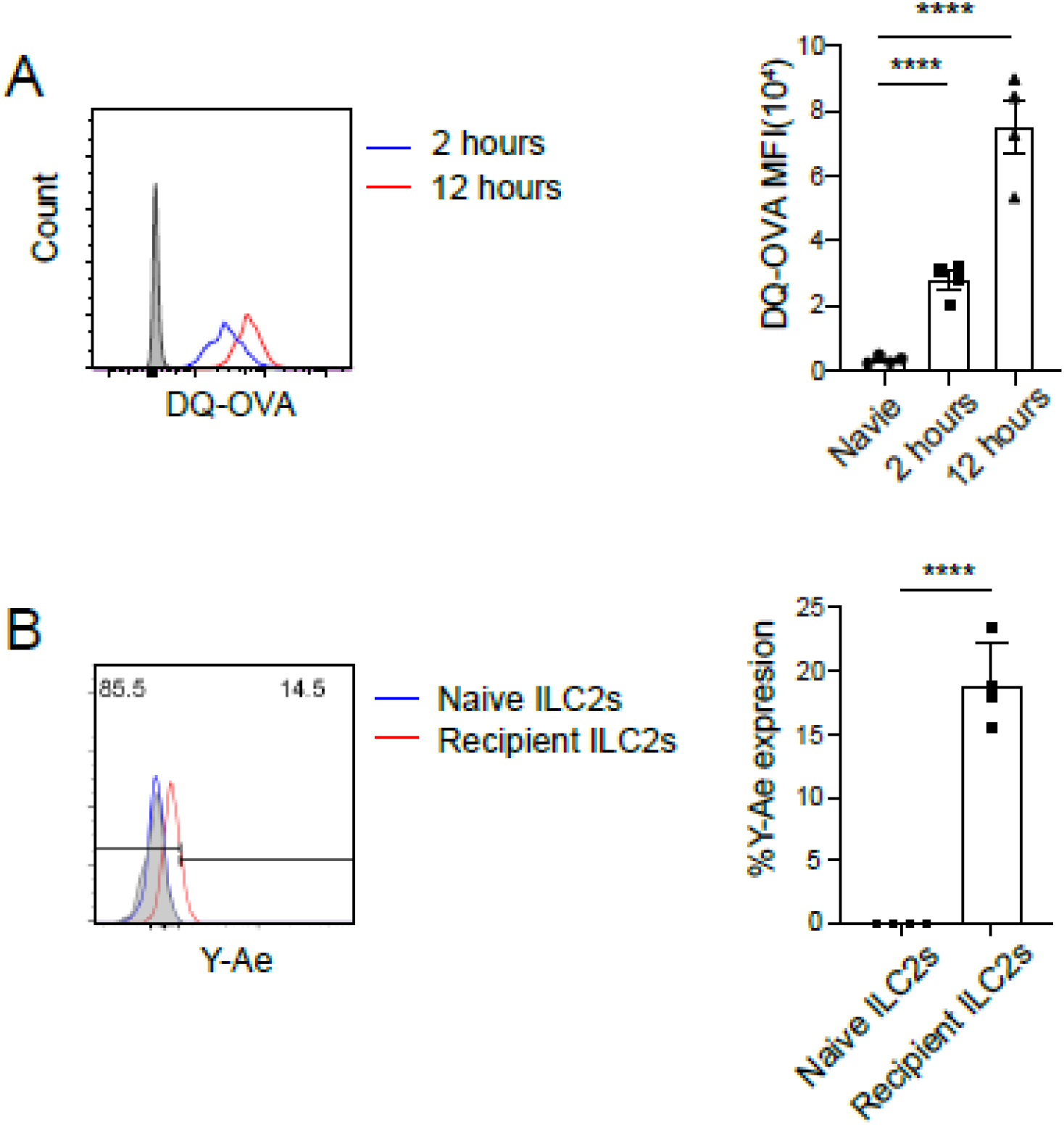
Infiltrating ILC2s can process and present alloantigen post allograft transplant. (A) Heart ILC2s were cultured with DQ-OVA and assayed for hydrolase induced fluorescence via flow cytometry in ILC2s. Left: Representative flow plot showing DQ-OVA signal expression on ILC2s after being cultured for indicated time with. ILC2s without DQ-OVA co-culture (grey) was used as negative control. Right: Quantitative analysis of the Mean Fluorescence Intensity (MFI) of DQ-OVA expression in ILC2s. (B) ILC2s from naive hearts and cardiac allografts at day 7 post mHTx and assayed via flow cytometry for Y-Ae expression. Left: Representative flow plot of Y-Ae on ILC2s from naive or heart allograft. FMO (fluorescence minus one, grey) was adopted as negative control. Right: Quantitation of Y-Ae in naïve vs heart allograft ILC2s. (n=4/group, data are presented as mean ± SEM; one-way ANOVA, **** p< 0.0001).

### Heart ILC2s induce CD4 T cell anergy in vitro due to the lack of co-stimulatory ligands

Given that infiltrating donor ILC2s can upregulate MHCII, process antigen, and present organ derived antigens in the absence of CD80/86, we assessed if ILC2s can also induce antigen specific CD4^+^ T anergy. To study this, we cultured heart derived Lin-CD127^+^ST2^+^ ILC2s with naïve OT-II T cells and soluble peptide (OVA_323-339_ peptide). Not surprisingly, ILC2s failed to induce the proliferation of OT-II in the presence of OVA peptide (Figure 6A and 6B). This contrasts with lung derived ILC2s, which express co-stimulation molecules constitutively, do activate antigen specific OT-II CD4^+^ T cells in the same setting (Supplemental Figure 1 and Figure 6C). When performing the proliferation assay, we also noticed that fewer OT-II T cells were retrieved after culturing in the presence of heart ILC2s compared to lung ILC2s (Figure 6D). To further clarify the role of MHCII on infiltrating cardiac ILC2s, we assayed for Caspase-3, known to be associated with T cell apoptosis following anergy^16^. Caspase 3 expression was elevated in graft infiltrating T cells in vitro. Further, blockade of MHCII partially rescued the loss of CD4s and decreased CD4 T cell Caspase-3 expression, suggesting that apoptosis is mediated by ILC2 antigen presentation (Figure 6E). LAG-3 expressed on CD4^+^ T cells can interact with MHCII from APCs, functioning as an inhibitory signal^17^. We indeed observed LAG-3 expression on OTII cells after transplant (Supplemental Figure 3A). To exclude the possibility that ILC2s suppress CD4^+^ T cell function through MHCII-LAG-3 interaction, we co-cultured OT-II cells with heart-derived ILC2s and OVA peptide in the presence of either anti-LAG-3 or an isotype control. As demonstrated in Supplemental Figure 3B, LAG-3 blockade did not affect OT-II cell numbers when cultured with ILC2s, indicating that the ILC2 suppression on CD4^+^ T cells was primarily due to ILC2 MHCII binding TCR rather than LAG-3 in our model. Taken together, these data suggest that ILC2s induce alloimmune T cell anergy in vitro via MHCII mediated antigen presentation, in the absence of CD80/86.

**FIGURE 6.**
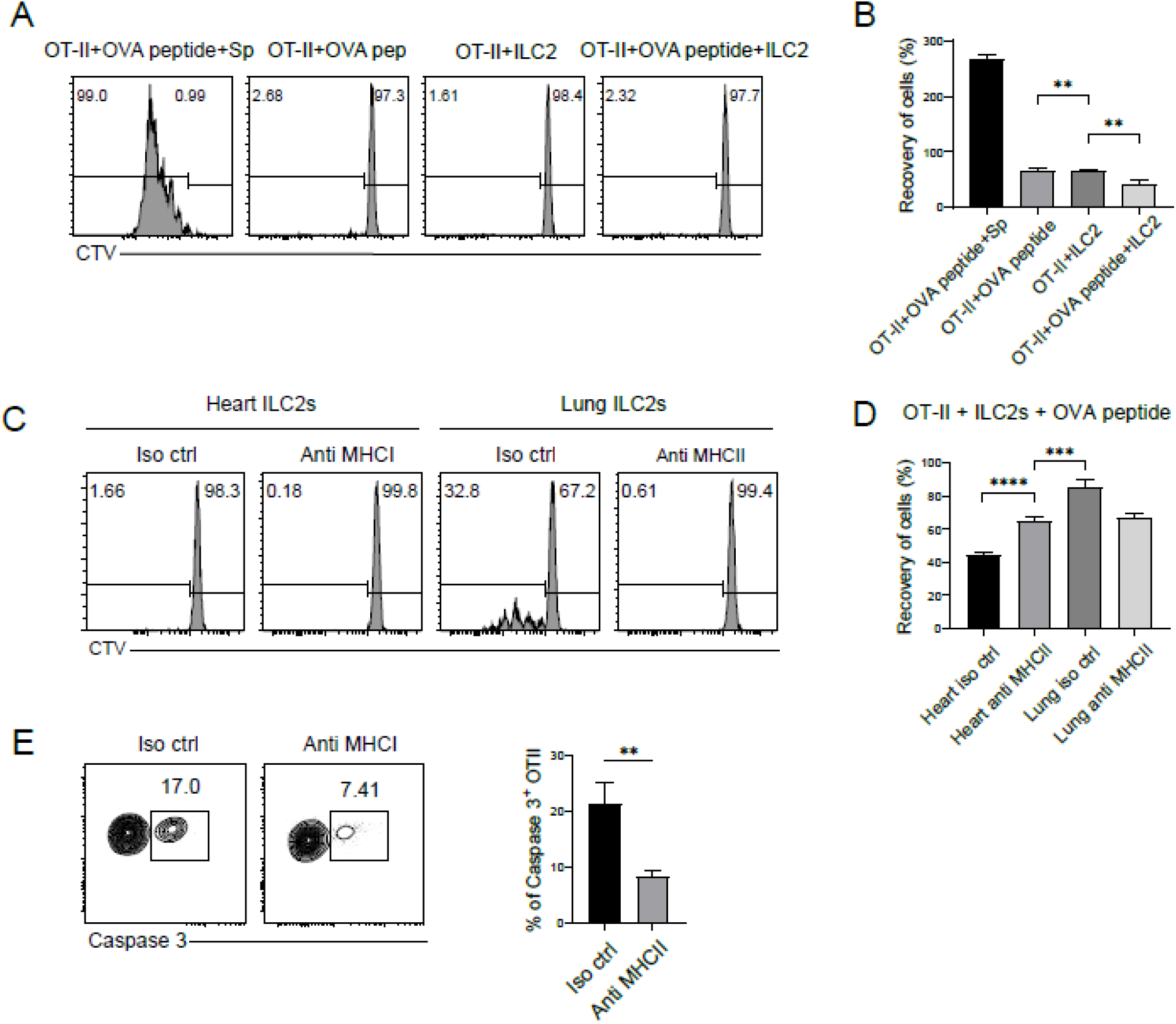
Heart ILC2s induce CD4^+^ T cell anergy and apoptosis in vitro. (A) Representative flow cytometry histograms showing CTV dilution of OT-II T cells cultured under different conditions. (B) Quantification of OT-II T cell proliferation under different culture conditions. **p<0.01. (C) Representative flow cytometry histograms comparing CTV dilution of OT-II T cells cultured with heart ILC2s or lung ILC2s, with or without anti-MHCII blocking antibody. (D) Quantification of retrieved OT-II T cell after culture with heart or lung ILC2s, with or without anti-MHCII blocking antibody. (E) Left: Representative flow cytometry plots showing Caspase-3 expression in OT-II T cells cultured with heart ILC2s, with or without anti-MHCII blocking antibody. Right: Quantification of Caspase-3+ OT-II T cells after culture with ILC2s. Data presented as mean ± SEM. **p<0.01, ***p<0.001, ****p<0.0001. Statistical analysis was performed using one-way ANOVA or unpaired two-tailed Student’s t-test.

### Heart ILC2s induce anergic alloreactive CD4^+^ T cell in heart grafts

To better understand the effect of heart ILC2s on donor specific CD4^+^T cells in vivo, we transplanted allogeneic OVA (B6 background) x Balb/C F1 hearts into either Rag2 KO (No T, B cells) or Rag2/Il2rg DKO (No T, B cells, NK / ILC2s). In allogeneic transplants, NK cells can be activated through “missing-self” mechanism (refers to NK cells recognized donor cells when they lack self-MHC class I molecules, leading to NK-mediated rejection)18. Therefore, B6 OVA × Balb/C donor hearts were adopted to eliminate NK cell-mediated effects. Anti-CD90.2 antibody was also given to donors prior to transplant to delete donor ILC2s and to ensure that donor ILC2s have no effect on our observations (Supplemental Figure 4). OVA specific OT-II transgenic CD4^+^ T cells were then adoptively transferred 10 days after transplant to ensure an adequate recovery of cardiac ILC2s, functioning as donor reactive T cells (Figure 7A). Four days after adoptive transfer, we observed more OT-II CD4 T in the heart grafts from Rag2/Il2rg DKO recipients compared to Rag2 KO recipients (Figure 7B-C). In addition, most of OT-II T cells in the hearts graft showed a CD44^hi^CD62L^lo^ effector T cell or CD44^hi^CD62L^hi^ memory phenotype, suggesting they had been activated within the allograft (Figure 7B). Interestingly, no naïve CD4^+^ T cells were induced to Treg cells (Supplemental Figure 5). These data suggest that the presence of ILC2s is able to limit the intragraft antigen specific T cells. Although both recipient groups produced similar amounts of IL2, OT-II T cells from Rag2 KO recipients displayed significantly less Ki67 expression, produced decreased amount of IFN-γ, suggestive of an anergic phenotype (Figure 7C).

**FIGURE 7.**
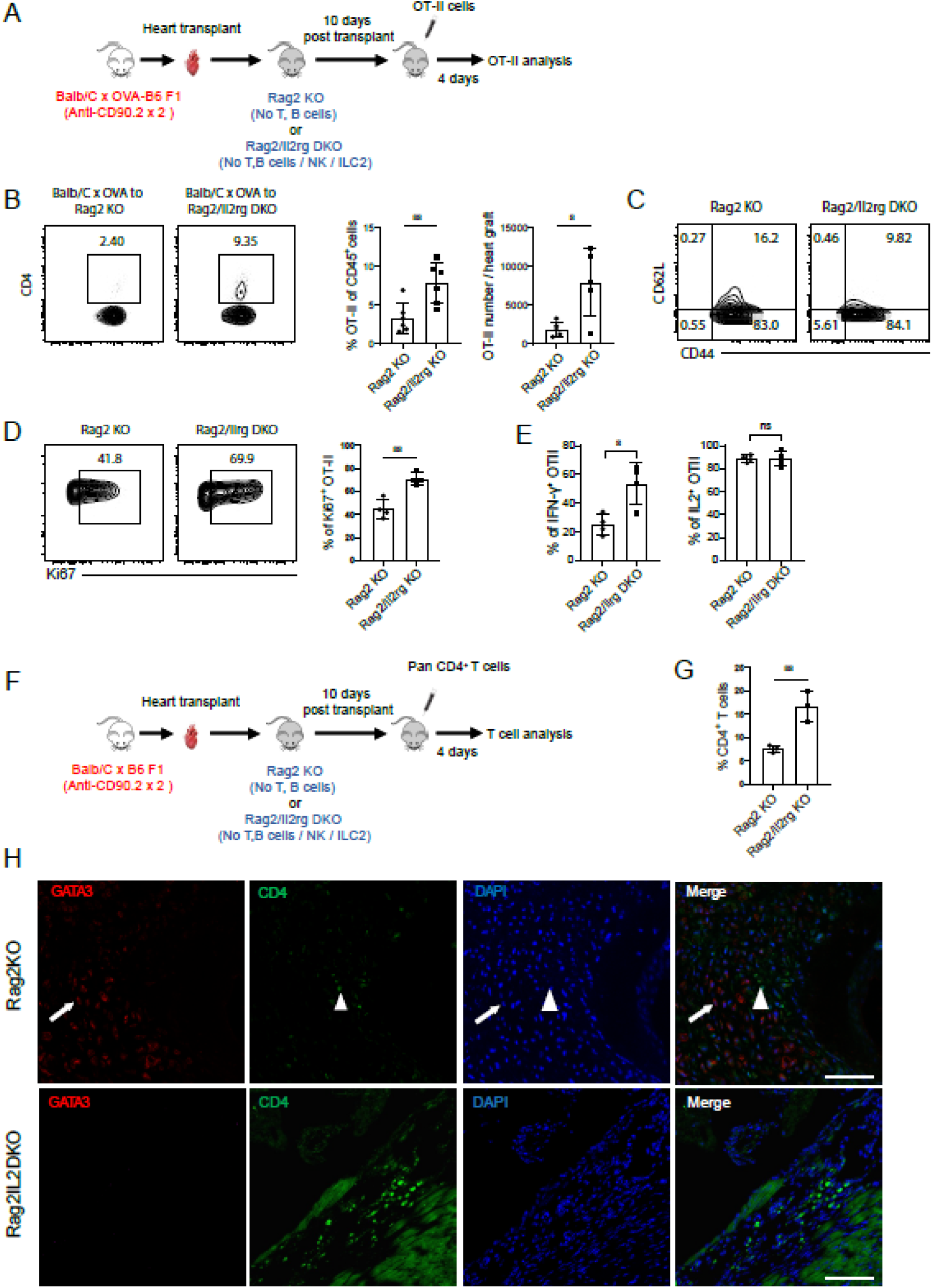
Infiltrating ILC2s prevent alloimmune T cells activation following solid organ transplant. (A) Experimental design: OVA (B6 background) x Balb/C F1 hearts were transplanted into either Rag2 KO (No T, B cells) or Rag2/Il2rg DKO (No T, B cells, NK / ILC2s) recipients and treated as described. (B) Flow cytometry analysis of OT-II cells in heart grafts. Left: Representative plots showing CD4^+^ OT-II cells. Middle: OT-II cell percentage among CD45+ cells. Right: Total number of OT-II cells per heart graft. Data are presented as mean ± SEM (n=4-6/group). Statistical analysis was performed using unpaired two-tailed Student’s t-test (*p<0.05, **p<0.01). (C) Representative plots showing CD44 and CD62L expression on OT-II cells from Rag2 KO and Rag2/Il2rg DKO recipients. (D) Proliferation of OT-II cells. Left: Representative plots of Ki67 expression in OT-II cells. Right: Quantification of Ki67+ OT-II cells. (E) Cytokine production in OT-II cells. Left: Percentage of IFN-γ+ OT-II cells. Right: Percentage of IL-2^+^ OT-II cells. Data are presented as mean ± SEM (n=4-6/group). Statistical analysis was performed using unpaired two-tailed Student’s t-test (*p<0.05, **p<0.01, ns: not significant). (F) Experimental design: Balb/C x B6 F1 hearts were transplanted into Rag2 KO or Rag2/Il2rg DKO recipients and treated as described. (G) Quantification of CD4^+^ T cells in heart grafts of Rag2 KO and Rag2/Il2rg DKO recipients. Data are presented as mean ± SEM (n=3/group). Statistical analysis was performed using unpaired two-tailed Student’s t-test (**p<0.01). (H) immunofluorescence microscopy (red= GATA3 positive cells; green = CD4, blue = DAPI and merged filter).

Since the OT-II targeting OVA system represents a single-antigen cognate allorecognition model, we also sought to determine whether ILC2s could similarly restrain pan alloimmune T cells in a similar setting. To study this, F1 progeny of a Balb/C and B6 cross were used as cardiac donors and transplanted into either Rag2 KO or Rag2/Il2rg DKO mice. Anti-CD90.2 antibody was again given to donors prior to transplant to delete donor ILC2s. Magnetically sorted B6 pan CD4^+^ T cells were then adoptively transferred 10 days after transplant (Figure 7D). Similar to our data in Figure 7B, we observe significantly fewer cardiac allograft CD4 T cells in Rag2 KOs, or in the presence of ILC2s (Figure 7E and F).

### Expansion of ILC2 in transplanted murine hearts leads to prolonged allograft survival

Our data prove that ILC2s upregulate MHCII in the absence of CD80/86, induce T cell anergy and apoptosis via Caspase-3 in vitro, restraining the activation of alloimmune CD4^+^ T cells within the transplanted organ. To further define the importance of ILC2s in suppressing the alloimmune response, we examined the survival of heart allografts in the presence or absence of ILC2s. No differences in survival were observed when donor-specific OT-II T cells targeting the minor antigen OVA were transferred (Supplemental Figure 5). However, with pan CD4^+^ T cells recognizing significantly more donor antigens in CB6F1, including murine major histocompatibility complexes, heart grafts were gradually rejected overtime. However, ILC2s significantly prolonged allograft survival (Figure 8A and 8B). Importantly, these differences in survival were preserved when NK cells were specifically depleted in Rag2 KO mice (Figure 8B and Supplemental Figure 6).

**FIGURE 8.**
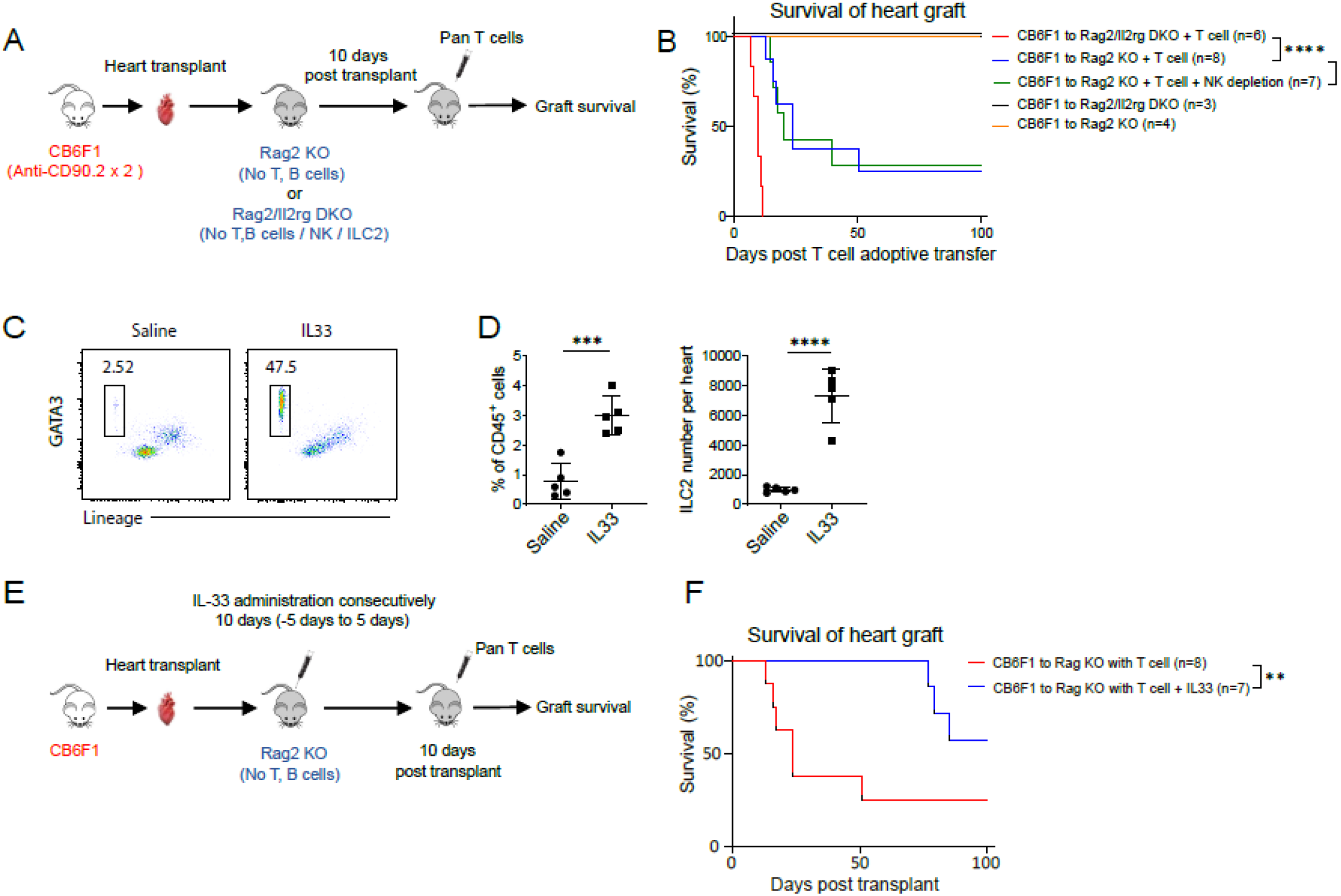
Expansion of ILC2s by IL33 administration In vivo promote allograft survival in murine heart transplantation. (A) Experimental schema. CB6F1 hearts (treated with anti-CD90.2) were transplanted into Rag2 KO or Rag2/Il2rg DKO recipients. Pan CD4^+^ T cells were adoptively transferred 10 days post-transplant. Graft survival was monitored. (B) Survival of allografts with and without T cell adoptive transfer in Rag2 KO vs Rag2/IL2rg DKO recipients accepting CB6F1 hearts, NK cells were depleted before transplant where indicated. (C) Representative flow cytometry plot illustrating IL-33 induced expansion of ILC2s in vivo. Cells isolated from hearts of saline or IL-33 administrated B6 mice were gated on CD45^+^ population. (D) Quantification of ILC2 expansion after IL-33 treatment. Left: Percentage of ILC2s among CD45^+^ cells. Right: Total number of ILC2s per heart (n=5/group, ***p<0.001, ****p<0.0001. Unpaired t-test). (E) Experimental schema of ILC2 expansion strategy. IL-33 was administered to Rag2 KO recipients for 10 consecutive days (−5 to ^+^5 days relative to transplant). Pan T cells were transferred 10 days post-transplant. (F) Survival curve showing the survival of mice following heart transplantation with or without ILC2 expansion by IL-33. For (B and F), Mice were monitored for 100 days post-transplantation. Statistical differences in survival between groups were determined using the Log-rank (Mantel-Cox) test. Data are presented as the percentage of survival over time (n=3-8/group. **** p<0.001, ** p<0.01, not significant).

It has been reported that IL-33 expanded kidney resident IL-10 producing ILC2s protected the islet allografts when being transplanted underneath the renal capsule and ILC2s infusion mitigate GVHD in GI tract. In contrast to these reports, in our model, IL-33 did not expand IL-10 producing heart ILC2s (Supplemental Figure 7). However, exogenous IL-33 administration expanded the ILC2s in heart tissue (Figure 8C and 8D). To test whether IL-33 treatment prolongs the survival of heart allografts, we conditioned recipient Rag2 KOs with and without IL-33 before and after mHTx (Figure 8E). IL-33 administration significantly increased the allograft survival (Figure 8F). These data highlight a novel role for ILC2s in constraining alloimmune CD4^+^ T cell responses through the induction of anergy and apoptosis, as well as demonstrate that ILC2s can be therapeutically manipulated through IL-33 administration to improve their effector function and allograft survival.

## Discussion

Over the course of the last decade, the role of innate lymphoid cells in disease has expanded dramatically. Group 2 innate lymphoid cells or ILC2s have been shown to participate in wound healing through the production of amphiregulin and collagen production, dampen ischemia reperfusion injury through the modulation of inflammatory responses via production of Th2 type cytokines, and have been shown to directly and indirectly communicate with CD4^+^ T cells^4,12,19^. In addition, as we and others have demonstrated, they are present in every tissue that we currently transplant, including heart, lung, kidney, and intestine^4,8^. Although these data support the possible beneficial function of ILC2s following solid organ transplantation, the role of ILC2s in alloimmunity remains unclear.

Our findings show that following murine cardiac transplantation, donor ILC2s are replaced by phenotypically different recipient ILC2s. Further, these migrating recipient ILC2s express MHCII in the absence of CD80/86 and can process and present self-antigen. We also demonstrate that in vitro coculture of cardiac allograft derived ILC2s loaded with OVA peptide can induce CD4^+^ T cell anergy. In addition, we also show a significant decrease in allograft infiltrating T cells and significantly improved allograft survival in the presence of ILC2s compared to mice without ILC2s. Taken together, these data suggest that migrating recipient ILC2s constrain the infiltration of alloreactive T cells, induce donor specific CD4^+^ T cell anergy, and facilitate allograft tolerance.

While there are limited studies investigating the role of ILC2s in transplant, these studies have also suggested a beneficial immunoregulatory role for ILC2s. For example, Huang et al demonstrated that administration of IL-33, expanded ILC2s which were at least partially protective against alloislet transplantation^20^. In addition, Huang et al also showed that 2 separate phenotypes of ILC2s expanded in response to IL-33, an IL-10 producing and a non-IL-10 producing group^20^. In addition, it was only the IL-10 producing ILC2 subset that provided alloislet protection in their model. While ILC2s are also protective in our cardiac allograft model, migrating recipient ILC2s into the cardiac allograft did not produce IL-10 either in situ or ex vivo when stimulated with IL-33 (Supplemental Figure 8).

Guo et al. also observed a beneficial role for ILC2s in transplant^19^. Using a murine lung allograft model, they showed that IL-33-dependent infiltration of ILC2s into the transplanted lung led to the production of significant amounts of IL-5, which subsequently recruited eosinophils, providing protection against alloimmunity^19^. Conversely, in our model, IL-5 expression decreases overtime in migrating cardiac allograft ILC2s suggesting that the mechanism for which ILC2s exert their immunoregulatory role following solid organ transplant may be tissue specific (Supplemental Figure 8). Certainly, our data and data from Shaikh et al. suggests that ILC2s can alter their phenotype depending on what tissue they reside in^19^.

In contrast to the above studies, the mechanism of ILC2 dependent allograft protection and induction of tolerance in our model is through ILC2-CD4^+^ T cell interaction and induction of T cell anergy. Other investigators have provided evidence of direct interaction between CD4^+^ T cells and ILC2. Oliphant et al. demonstrated the upregulation of MHCII on ILC2s and direct CD4-ILC2 interaction which resulted in the activation of Th2 anti-helminth immunity^12^. Mirchandani et al. also documented that expanded lung ILC2s can stimulate CD4^+^ T cell proliferation and enhance Th2 dependent allergen responses21. Interestingly, both lung and mesenteric lymph nodes ILC2s express CD80/86 and so as opposed to the T cell anergy that we observe in our study, these prior studies demonstrate CD4 proliferation in ILC2-CD4 coculture experiments.

The question of why ILC2s are not enough to completely constrain alloimmunity and contribute to tolerance in immunocompetent models of mHTx or clinically is unclear. It’s possible that their migration into newly transplanted allograft is impaired by interferon gamma (IFNγ) produced during the alloimmune response. Cautivo et al. recently demonstrated that interferon gamma limited the parenchymal distribution of ILC2s following allergic type inflammation to the adventitia of vessels^9^. While the inflammation induced through solid organ transplant is likely different, blockade of IFNγ could be employed as a strategy to improve ILC2 migration following solid organ transplant. However, IFNγ has also been found to be important for the development of T regulatory T cells (Tregs) which are known to be important for the long-lasting peripheral tolerance^22^. So cell specific and not systemic blockade of IFNγ would be important. It is also possible that because ILC2s have to migrate from their niche and likely phenotypically switch to a cardiac specific ILC2 at the same time alloimmune cells are infiltrating into the newly transplanted solid organ, there is insufficient time to perform their immunoregulatory role. Therefore, strategies to enhance their presence within the allograft and increase their immunoregulatory role would be critically important to improve their impact on allograft tolerance.

Our studies reveal a novel role for ILC2s in the inhibition of alloimmunity and induction of antigen specific CD4^+^T cell anergy following solid organ transplant. Further, while other studies have demonstrated that ILC2s have an immunoregulatory role in murine lung and islet transplant through either a paracrine effect from suppressive cytokines or through stimulation of Th2 dependent responses, this is the first study to show evidence that ILC2s can induce T cell anergy in transplantation ^19–21^. Regardless, these published findings do not take away from the importance of our results and further suggest that targeting ILC2s to augment their responses post-transplant would be an important adjunct to our current therapeutic strategies. Considering that ILC2s are tissue resident, present in every organ that is currently transplanted, and have been proved to engage in cross talk with other immunoregulatory populations, it is possible that ILC2s are the first interface between the newly transplanted organ and host immunity. Further, as we have shown that ILC2s also facilitate tolerance, strategies to exploit the immunoregulatory function of ILC2s to improve outcomes in solid organ transplantation should be explored.

## Materials and methods

### Mice

B6 (H2-Kb), B6 pep-boy (CD45.1), Balb/C (H2-Kd), DBA/2J (H2-Kd), Rag2 KO (B6 background), Rag2/Il2rg DKO (Balb/C background) and OT-II TCR transgenic mice were purchased from the Jackson Laboratory. Rag2/Il2rg DKO (B6 background) mice were obtained from Taconic. Eight to 12 weeks old male mice were used in this study and maintained under specific pathogen-free conditions. All animal work was performed in compliance with ethical regulations and was approved by the Institutional Animal Care and Use Committee (IACUC) of the Boston Children’s Hospital.

### Mouse surgery procedure and drug administration

Heterotopic heart transplantation was performed as described previously^23^. Briefly, heart grafts were harvested from donors and transplanted into the abdominal cavity of recipient mice by anastomosing the donor aorta and pulmonary artery to the recipient’s aorta and vena cava respectively in an end to side manner. Graft survival was monitored by palpation and rejection was defined as cessation of heartbeat, further verified by laparotomy. For in vivo depletion of NK1.1 expressing cells, mice were injected i.p every day for post-operative 7 days with 25μg of InVivoMAb anti mouse NK1.1 (clone: PK136 Bio X Cell). A carrier-free, recombinant mouse IL-33 (0.5ug/day, Biolegend) was administered to each recipient intraperitoneally from 5 days before transplantation until 5 days post-transplantation.

### Tissue digestion and cell isolation

Single cell suspensions were prepared from tissues including native hearts, transplanted heart grafts, lungs, mesenteric lymph nodes (mLN) and spleens. For heart and lung disassociation, perfusion was performed through inferior vena cava with heparin saline (50U/L) and digestion cocktail consisting of HBSS, collagenase A (1mg/ml) and DNAse (0.2mg/ml). Then samples were minced into small pieces after being excised and digested at 37°C for 60 min in shaking incubator. The remaining tissue was mashed with a syringe plunger and single cell suspension was filtered through 40 μm cell strainer. For splenocytes and mLN cells isolation, tissues were directly mashed through cell strainers without enzyme digestion. Single cell suspension was subsequently subjected to density gradient centrifugation with Ficoll solution to enrich lymphocytes. Purified lymphocytes were used for analysis or further isolation through FACS sorting.

### Flow cytometry analysis and cell sorting

Single cell suspensions were incubated on ice with anti-CD16/32 antibody for FC receptor block for 5 min, followed by fluorochrome conjugated antibodies in FACS buffer (PBS with 2% FBS and 1mM EDTA). Dead cells were excluded using Fixable Viability Dye eFluor 780 (Thermon Fisher Scientific, Cat: 65-0865-14). For ILC2 staining, linage positive cells were excluded by staining with APC anti-mouse Lineage Cocktail (BD Biosciences, Cat: 558074). Surface marker staining was performed by incubating cells with antibodies on ice for 30 min. following antibodies were used for surface staining: CD45 (BioLegend, 11-0451-82, Clone: 30-F11), CD127 (BioLegend, Cat: 135037, Clone: A7R34), CD25 (BioLegend, Cat: 102033, Clone: PC61), ST2 (ThermoFisher, Cat: 12-9335-82, Clone: RMST2-2), CD90 (BioLegend, Cat: 53-2.1, Clone: 25-0902-81), KLRG1 (BioLegend, Cat: 138421, Clone: 2F1/KLRG1), H2-Kb (BioLegend, Cat: 116510, Clone: AF6-88.5), H2-Kd (BioLegend, Cat: 116620, Clone: SF1-1.1), CD45.1 (BioLegend, Cat: 110717, Clone: A20), MHCII (BioLegend, Cat: 107615, Clone: M5/114.15.2), I-Ad (BioLegend, Cat: 115008, Clone: 39-10-8), Y-Ae (Thermo Fisher, Cat: 11-5741-82, eBioY-Ae), CD80 (BioLegend, Cat: 104725, Clone; 16-10A1), CD86 (BioLegend, Cat: 105123, Clone: PO3), CD44 (BioLegend, Cat: 103010, Clone: IM7), CD62L (BioLegend, Cat: 104412, Clone: MEL-14), LAG-3 (BioLegend, Cat: 125210, Clone: C9B7W). Samples were fixed by 4% PFA before data collection when necessary. For intracellular protein labelling, cells were permeabilized and fixed with Transcription Factor Staining Kit (Invitrogen, Cat: 00-5123-43). When cytokine staining was required, isolated cells were cultured with Cell Stimulation Cocktail (ThermoFisher, Cat: 53-4875-82, Clone: Dan11mag) and GolgiPlug (BD Biosciences, Cat:555029) at 37°C for 5 hours. Following antibodies were cultured with cells at 37°C for intracellular markers: GATA3 (BioLegend, Cat: 653804, Clone: 16E10A23), Eomes (BioLegend), T-bet (BioLegend, Cat: 644803, Clone: 4B10), RORγT (BD, Biosciences, Cat: 562894, Clone Q31-378), IL-5 (BioLegend, Cat: 504311, Clone: TRFK5), IL-13 (BioLegend, Cat: 159408, Clone: W17010B), IL-10 (BioLegend, Cat: 505021, Clone: JES5-16E3), IL-9 (BioLegend, Cat: 514109, Clone: RM9A4), IFN-γ (BioLegend, Cat: 505815, Clone: XMG1.2), IL-2 (BioLegend, Cat: 503825, Clone: JES5-5H4), Ki67 (BioLegend, Cat: 652405, Clone: 16A8), Foxp3 (BioLegend, Cat: 126408, Clone: MF-14). Precision counting beads (BioLegend, Cat: 424902) were used for ILC2s absolute number assessment. All flow cytometry experiment data were acquired using a BD LSR Fortessa flow cytometer with FACS Diva software (BD Biosciences) and analyzed with FlowJo V10.8.1.

ILC2s purification was performed through FACS sorting. Single cell suspensions obtained from tissues were stained with CD45, Lineage cocktail, CD127 and ST2. Lin-CD45+ CD127+ ST2+ population were identified as ILC2s. Cells were sorted by BD FACSAria II. Antibodies used in flow cytometry and sorting experiments were purchased from eBioscience, BioLegend or BD Biosciences if not otherwise indicated.

### Bulk RNA sequence of ILC2

Total RNA was extracted from allograft or naïve heart ILC2s using the Universal RNA Extraction CZ Kit (RNC643, ONREW) according to the manufacturer’s instructions. RNA quantity was analyzed using Qubit 4.0 (Invitrogen) and quality was examined by electrophoresis on a denaturing agarose gel. Total RNA was used as input material. cDNA synthesis and amplification were performed using the Single Cell Full Length mRNA-Amplification Kit (N712-03, Vazyme), and purification was done with VAHTS DNA Clean Beads (N411-02, Vazyme). Purified cDNA quality was examined by gel electrophoresis and quantified using Qubit 4.0 (Invitrogen). Libraries were prepared according to the TruePrep® DNA Library Prep Kit V2 (TD502-02, Vazyme) user manual. Sequencing was performed using the Illumina Novaseq 6000 platform with a 150 bp paired-end strategy. Differentially expressed genes (DEGs) between 4 allograft ILC2 and 3 naive B6 heart ILC2 samples were identified using the DESeq2 (v 1.32.0) package (https://pubmed.ncbi.nlm.nih.gov/25516281/) (P < 0.05 and |log2-fold change (FC)| > 0.585). Volcano plots and heatmaps were generated using the ggplot2 (v 3.3.2) (https://pubmed.ncbi.nlm.nih.gov/24132163/) and heatmap (v 0.7.7) packages, respectively. Gene Ontology (GO), Hallmark, and Kyoto Encyclopedia of Genes and Genomes (KEGG) pathway analyses of the DEGs were performed with the clusterProfiler (v 3.16.0) (https://pubmed.ncbi.nlm.nih.gov/34557778/) and enrichR (v 3.2.0) (https://pubmed.ncbi.nlm.nih.gov/27141961/) packages (P < 0.05). All data was upload to GEO (GSE290285).

### ILC2s culture and immune assay

Before ILC2s isolation, mice were administered of recombinant IL-33 (Biolegend) as detailed in “antibodies and cytokines administration” section. Sorted ILC2s from hearts or lungs were immediately plated in complete medium (RPMI1640 with 10% FBS, 200U/ml Pen/Strep, sodium pyruvate, MEM non-essential amino acids and 2-ME) supplemented with 10ng/ml IL-7 (Thermo Fisher Scientific) and 10ng/ml IL-33. Cells were cultured in 37 °C and 5% CO2 and mediums were changed every 3 days. ILC2s were maintained in vitro for 6 days. Cell purity was confirmed by flow cytometry by the end of culture. Where indicated, 1.5×10^4^ ILC2s were cultured with 10 μg/ml DQ-OVA (Thermo Fisher Scientific) at 37 °C for 2 hours or 12 hours. Cells were washed intensively with FACS buffer and fluorescence was immediately assessed by flow cytometry.

### OT-II CD4^+^ T cells proliferation assay

Mouse Naïve CD4^+^ T cells were isolated from OT-II TCR transgenic mice spleens using EasySep mouse naïve CD4^+^ T cell isolation kit according to manufacturer’s protocol. 2×10^4^ ILC2s were then cultured with 2×10^4^ naïve OT-II T cells in complete medium supplemented with 1ng/ml rIL-2 (Sigma Aldrich) at 37 °C in 5% CO2 incubator. 4×10^4^ splenocytes from naïve B6 mice or soluble OVA323-339 peptide (10 μg/ml) were added into culturing as stated. To block the MHCII-TCR interaction, anti MHCII antibody (clone M5/114.15.2,) or isotype control were used at a concentration of 1 μg/ml. AntiLAG-3 antibody (clone C9B7W, Bioxcell) or Isotype control were used to block MHCII-LAG-3 interaction. OT-II T cells numbers were calculated by flow cytometry with precision counting beads by the end of 72 hours culturing.

### Immunofluorescence analysis

Heart grafts were fixed with 4% paraformaldehyde in room temperature for 1 hour. Then fixed tissues were washed in PBS thoroughly and dehydrated in 30% sucrose solution in 4°C overnight. After embedding in OCT Compound (Sakura Finetech), vertical cryosections were made at 10 microns and stored at –20°C for storage. The sections were rehydrated with PBS and then blocked and permeabilized for 1 hour with 10% normal goat serum (50062Z, Thermo Fisher), 1% bovine serum albumin (Sigma-Aldrich) and 0.1%Triton-X100. For biotinylated primary antibody, samples were further blocked using an Endogenous Biotin-Blocking Kit (E21390, Invitrogen) according to manufacturer’s instructions. Slices were incubated with primary antibody at room temperature for 1 hour in blocking solution. Subsequently, they were washed with PBS and incubated with secondary at room temperature for 1 hour. Slides were mounted in DAPI Fluoromount-G medium (Southern Biotech). Images were taken by Zeiss LSM880 confocal scanning microscope. The secondary antibodies were listed as followed: Goat anti-Rabbit IgG H&L highly cross-adsorbed secondary antibody (1:500 for section staining, 1:200 for whole mount staining, Alexa Fluor Plus 647, A32733, Thermo Fisher), Goat anti-Rabbit IgG H&L (1:500 for section staining, 1:200 for whole mount staining, Alexa Fluor 488, ab150077, Abcam), Streptavidin (1:500, Alexa Fluor 647, S32357, Thermo Fisher).

## Acknowledgements

Funding: AGC is supported by the Hardy Hendren Faculty Development fund, the Boston Children’s Translational Research Program Junior Translational Investigator Science award, and the BCH OFD BTREC-CRTREC faculty development award.

## Author contributions

JG, WP, and ZX designed and performed experiments, analyzed data and contributed to manuscript writing/editing. TB contributed to data analysis and experimental design. IZ contributed to manuscript writing and editing. AGC designed experiments, contributed to data analysis, and manuscript writing/editing.

## Data and Materials availability

Sequencing data have been depositing to GEO under the accession number GSE290285. All other data needed to evaluate the conclusion in this manuscript are included in the paper or supplemental materials.

