## Supplemental File for "Group 2 Innate lymphoid cells promote allograft survival by constraining and inducing anergy in alloreactive CD4^+^ T cells"

**SUPPLEMENTAL FIGURES AND MATERIALS**

**Supplemental Figure** **1. Distinct CD80/CD86 expression on ILC2s across different tissue types.** Representative flow cytometry scatter plots depict CD80 and CD86 on naïve or IL-33 stimulated ILC2s from mesenteric lymph nodes (mLN), kidney and lung. Left: Gated on Cd45^+^ cells from tissue. Middle: Gated on lineage^-^GATA3^+^CD127^+^ ILC2s. Right: Plots of CD80 and CD86. Fluorescence minus-one (FMO) samples were used to determine the gating strategy.

**Supplemental Figure 2**. **No cross dressing was found in immunocompromised recipients.**

Representative histogram showing donor MHCII (I-Ad) expression on recipient ILC2s (H2-Kb^+^) isolated from heart allografts 21 days after Balb/C to Rag2 KO B6 transplant.

**Supplemental Figure** **3. The expression of LAG-3 on OT-II does not affect T cell function during interaction with ILC2 MHCII.** **(A)** Representative histogram graph of LAG-3 expression on OT-IIs isolated from spleen (red) and heart allograft (blue). OT-II cells were adoptively transferred into Rag2 KO recipient following (Balb/C x OVA B6) F1 heart graft implant, and retrieved 7 days after adoptive transfer for LAG-3 analysis. FMO (grey) sample was used to determine the gating strategy. **(B)** Bar graph showing recovered OT-II cells after being cultured with OVA peptide and heart ILC2s with or without anti LAG-3 blockage. Data are presented as mean ± standard error of the mean (SEM). Statistical significance was determined by unpaired t-test. p value < 0.05 was considered statistically significant (ns, not significant).

**Supplemental Figure** **4. ILC2 depletion by anti-CD90.2 antibody administration.** Flow cytometry scatter plots showing heart ILC2s from isotype (left) or anti-CD90.2 antibody (right) conditioned mice.

**Supplemental Figure** **5. Differentiation of Treg from OT-II upon adoptive transfer. (A)** Representative flow cytometry plots showing Foxp3^+^ Tregs induction (gated on CD4^+^ cells) from OT-II in heart graft and recipient spleen. Analysis was conducted 7 days after naïve OT-II T cells adoptive transfer into Rag2 KO or Rag2/IL2rg DKO recipients following (Balb/C x OVA B6) F1 heart transplant. (n=4-5/group). **(B)** Bar graph presidents the percentage of Foxp3^+^ Tregs in OT-II cells from spleen or heart grafts of Rag2 KO or Rag2/IL2rg DKO recipients. Data are presented as mean ± standard error of the mean (SEM). Statistical significance was determined using unpaired t-test. p value < 0.05 was considered statistically significant (ns, not significant).

**Supplemental Figure** **6. OT-II cells donor reactive T cells do not cause graft loss in minor antigen miss-matched heart transplant.** Kaplan-Meier survival curve showing the survival of heart grafts following transplantation. Mice were monitored for 100 days post-transplantation. Statistical differences in survival between groups were determined using the Log-rank (Mantel-Cox) test. Data are presented as the percentage of survival over time (ns, not significant).

**Supplemental Figure 7. NK cells depletion in Rag2 KO mice. (A)** Representative flow cytometry scatter plots illustrating the percentage of NK cells in the Rag2 KO mice or giving Anti NK1.1 antibody *in vivo* for 1 week to deplete NK cells in the Rag2 KO mice. **(B)** Bar graph illustrating the percentage of NK cells in naïve Rag2 KO heart and heart allografts (CB6F1 to Rag2 KO heart transplant) 7 days or 14 days post-transplant after giving anti NK1.1 antibody for 1 week. Data are presented as mean ± standard error of the mean (SEM). Statistical significance was determined using one-way ANOVA. p value < 0.05 was considered statistically significant. ****p < 0.0001.

**Supplemental Figure** **8. Heart ILC2s do not express IL-10 in stable or IL-33 activated state.** Representative flow cytometry scatter plots illustrating IL-10 production by naïve or IL-33 stimulated ILC2s from hearts of different mouse strains.

**Supplemental Figure** **9. Diminished IL-5 production by grafting infiltrating ILC2s after heart transplant.** **(A)** Representative flow cytometry scatter plots showing IL-5 and IL-13 produced by ILC2 from naïve Balb/C heart and heart allografts (Balb/C to B6 heart transplant) 3 days and 5 days post-transplant. Data are representative of 2 independent experiments (n=2-3/group). **(B)** Bar graph illustrating the percentage of IL-5^+^ or IL-13^+^ ILC2s in naïve Balb/C heart and heart allografts (Balb/C to B6 heart transplant) 3 days or 5 days post-transplant. Data are presented as mean ± standard error of the mean (SEM). Statistical significance was determined using unpaired t-test. p value < 0.05 was considered statistically significant. *p < 0.05, ** p < 0.01, ns: not significant.

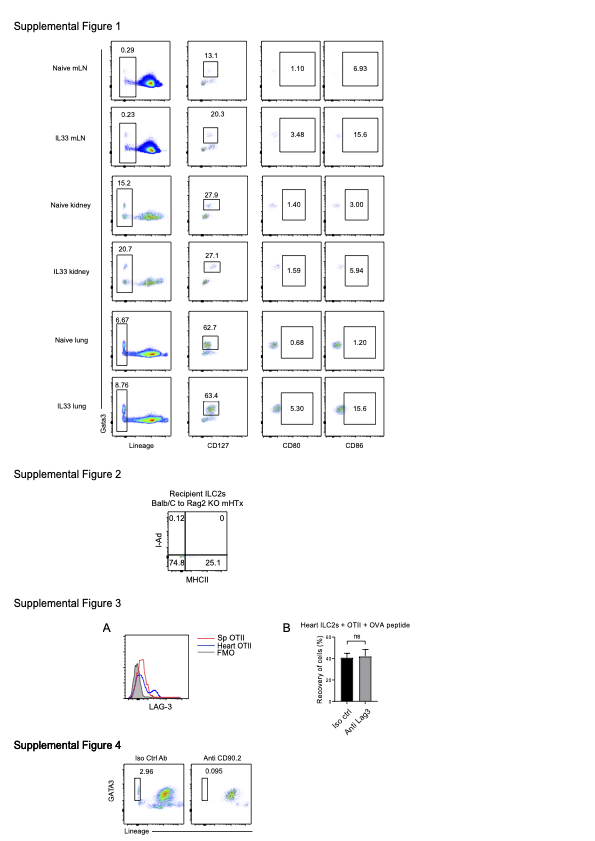

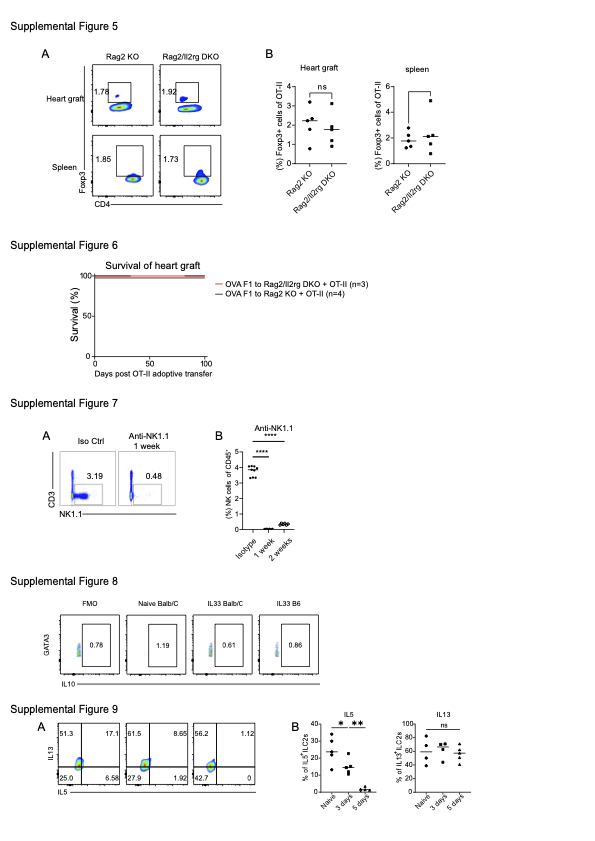
